# FBH1 deficiency sensitizes cells to WEE1 inhibition by promoting mitotic catastrophe

**DOI:** 10.1101/2023.05.15.540841

**Authors:** Lucy Jennings, Heather Andrews Walters, Jennifer M. Mason

## Abstract

WEE1 kinase phosphorylates CDK1 and CDK2 to regulate origin firing and mitotic entry. Inhibition of WEE1 has become an attractive target for cancer therapy due to the simultaneous induction of replication stress and inhibition of the G2/M checkpoint. WEE1 inhibition in cancer cells with high levels of replication stress results in induction of replication catastrophe and mitotic catastrophe. To increase potential as a single agent chemotherapeutic, a better understanding of genetic alterations that impact cellular responses to WEE1 inhibition is warranted. Here, we investigate the impact of loss of the helicase, FBH1, on the cellular response to WEE1 inhibition. FBH1-deficient cells have a reduction in ssDNA and double strand break signaling indicating FBH1 is required for induction of replication stress response in cells treated with WEE1 inhibitors. Despite the defect in the replication stress response, FBH1-deficiency sensitizes cells to WEE1 inhibition by increasing mitotic catastrophe. We propose loss of FBH1 is resulting in replication-associated damage that requires the WEE1-dependent G2 checkpoint for repair.

## 1. INTRODUCTION

Cancer cells often exhibit a high level of replication stress that can arise due to factors including oncogene expression and underlying DNA repair defects. Cells entering mitosis with high levels of replication damage undergo mitotic catastrophe and cell death [1]. To tolerate high levels of replication-associated damage, cancer cells become addicted to cell cycle checkpoints that arrest cells at the G2/M transition to allow cells time to repair. An emerging chemotherapeutic strategy is to target cell cycle checkpoints in cancer cells with high levels of replication stress to force premature entry into mitosis.

The WEE1 kinase controls CDK activity by phosphorylation of tyrosine 15 (Y15) on CDK1 and CDK2. WEE1 prevents entry of cells into mitosis by phosphorylation of CDK1 at tyrosine (Y15) arresting cells at the G2/M transition [2]. If DNA damage is not detected, phosphorylation of CDK1 on Y15 is removed by CDC25 phosphatase allowing cells to enter mitosis. At the beginning of S phase, CDK2 activity triggers firing of origins to initiate replication and this activity is controlled by WEE1 phosphorylation at Y15 [3–5]. WEE1 inhibition results in unscheduled origin firing resulting in a rapid depletion of nucleotide pools. The reduction in nucleotides results in accumulation of single stranded DNA leading to exhaustion of RPA pools and replication catastrophe [3,6]. Furthermore, WEE1 prevents cleavage of replication forks by the nuclease SLX4/MUS81 to produce double strand breaks and prevents nuclease-mediated degradation of stalled replication forks [3,5,7,8].

Given the roles of WEE1 in both regulation of S phase and the G2/M checkpoint, inhibition of WEE1 is an attractive chemotherapeutic target for cancer cells with high levels of replication stress. WEE1 inhibition kills tumor cells alone, but WEE1 inhibitors are much more effective in combination with inhibitors or treatments that increase replication stress such as those that target ATR and CHK1 kinases [9–11]. WEE1 inhibition alone kills cells with defects in the homologous recombination, the Fanconi anemia pathway, and may be more lethal to p53-deficient tumors [12–14]. These studies indicate that certain genetic defects sensitize cells to WEE1 inhibition highlighting a need for a better understanding of genetic alterations that modulate the response to WEE1 inhibitors.

The FBH1 helicase is critical for the induction of apoptosis in response to replication stress. FBH1 plays a role in replication fork reversal in response to hydroxyurea [15]. After prolonged replication stress, FBH1 cooperates with MUS81 to induce a double strand break at stalled forks to initiate apoptosis [16,17]. FBH1-deficient cells have decreased double strand break formation and increased survival in response to hydroxyurea and ultra-violet (UV) light. FBH1 has anti-cancer functions by protecting melanocytes from UV-mediated transformation. Hemizygous and homozygous loss of FBH1 has been identified in up to 63% of melanoma cell lines derived from metastatic melanoma[18].

Given the role of FBH1 in promoting MUS81-dependent cleavage of replication forks, we determined how FBH1-deficiency modulates the cellular response to WEE1 inhibition. As expected, we found FBH1 knockout (KO) cells results in less ssDNA accumulation accompanied a reduction in double break signaling indicating FBH1 is required for the efficient induction of the replication stress response due to WEE1 inhibition. Despite the reduction in replication catastrophe, we found that FBH1 KO cells are more sensitive to WEE1 inhibition due to a significant increase in pan-nuclear γH2AX and nuclear abnormalities consistent with mitotic catastrophe. We propose the underlying defect in the replication stress response in FBH1 KO cells is resulting in dependency of FBH1-deficient cells on the G2/M checkpoint to repair damage prior to mitosis. Our results suggest FBH1-deficient tumors may be sensitized to WEE1 inhibition.

## 2. MATERIAL AND METHODS

### 2.1. Cell lines and drug treatments

U2OS cells were grown in DMEM media (Gibco Cat #11965-092) +10% FBS (EqualFetal, Atlas Biologicals). Cell lines were cultured at 37°C, 5% CO2. Adovasterib (aka AZD1775; MedChemExpress) was resuspended in DMSO. Cells were treated with adovasterib as indicated.

### 2.2. Generation of FBH1 knockout U2OS cell line

We synthesized gRNAs flanking exon 4 and 5 and cloned them into the gRNA cloning vector (Addgene # 41824) using Gibson assembly (New England Biolabs)[19]. The targeting sequences for gRNAs were: FBH1 gRNA-1 5’-GCGGGGCACTTTGTGAGTGG-3’ and FBH1 gRNA-2 5’-GCGGCATGTTATTTCACTGG-3’. Cells were co-transfected with FBH1 gRNA-1, FBH1 gRNA-2, hCas9 (Addgene# 41815) and pcDNA 3.1 Hygromycin (Invitrogen) using Lipofectamine 3000 as per manufacturers’ instructions. Single colonies were isolated after selection in 100 ug/ml hygromycin. After growing colonies to a 96-well plate, clones were split into two 96-well plates. Genomic DNA was isolated from one well using QuickExtract solution following manufacturers’ protocol. Clones were screened for deletion of Exons 3 and 4 by PCR using Phusion polymerase. Primers used in screening: FBH1_F 5’-CCTGATGACCCCAAGCATG-3’, FBH1_R1 5’-GGTCCCGGTAGCCTGGTTAC-3’, and FBH1_R2, 5’-AAGATTCCTCAACCAAATGG-3’. Clones positive by PCR were expanded and screened for loss of FBH1 by Western blotting.

### 2.3. Cell titer blue assay

U2OS or FBH1-KO cells were plated in triplicate at various concentrations: 1000, 500, 250, and 125 cells in a 96-well plate. Cells were treated with either DMSO or 400 nM adovasterib for 48 hrs and allowed to outgrow for 4 or 5 days. Cells were stained with the CellTiter Blue reagent (Promega) as per manufacturers’ instructions. Fluorescence of each sample was read at an excitation of 570 nm and emission of 600 nm using Synergy H1 Plate reader (Biotek). The absorbance readings for each cell number (DMSO and adovasterib) were averaged. The percent survival was determined by dividing the average absorbance value of adovasterib-treated cells by the corresponding absorbance reading in untreated sample value. Experiment was performed in duplicate at least two independent times.

### 2.4. Immunofluorescence

U2OS or FBH1 KO (50,000) cells were plated on coverslips in a 12-well tissue culture plate and grown overnight. Cells were treated with various concentrations of adovasterib or DMSO for the indicated time points. Cells were incubated with HEPES/Triton X-100 buffer (20 mM HEPES, ph7.4, 0.5% TritonX-100, 50 mM NaCl, 3 mM MgCl2, 300 mM sucrose) for 10 min at room temperature. Cells were fixed with 3% paraformaldehyde, 3.4% sucrose in 1X PBS for 10 min at room temperature. Cells were washed with 1X PBS 3 times. Cells were blocked with 3% (w/v) bovine serum albumin (BSA) in PBS for at least 20 min at room temperature. Primary antibodies were diluted in 1% BSA and added to coverslips in a humidified glass chamber and incubated overnight at 4°C. Cells were washed 3 times with 1X PBS and then incubated with AlexaFluor-conjugated secondary antibodies (Invitrogen) in 1% BSA in a humidified chamber at room temperature for 1 hr. Cells were washed 3 times in 1X PBS. Cells were dehydrated by incubating for 2 min in increasing concentrations of EtOH (70%, 95%, and 100%) before mounting with Vectashield with DAPI (Vector laboratories). Images of at least 150 nuclei were acquired at 20X using a Zeiss Imager.M2 epifluorescence microscope equipped with a Axiocam 503 mono camera. Images were analyzed using Fiji software (version 1.53v, NIH) and data were graphed using Graphpad Prism software (version 9.5.0, Dotmatics).

### 2.5. Antibodies

Primary antibodies against γH2AX (ab26350, 1:1000), pH3 Ser10 antibody (ab5176, 1:1000), pKAP1 (ab70369, 1000), pCDK Y15 (ab275958, 1:1000), RPA (ab2175, 1:1000), KAP1 (ab22553, 1:1000), CDK1 (ab32094, 1:1000), and FBH1 (ab58881, 1:100) were from Abcam. BrdU antibodies (recognizes IdU) was from BD biosciences (347580, 1:50). pRPA S4/8 antibodies (A300-245A, 1:1000) was from Fortis Life Sciences. Tubulin antibodies was from Novus Biologicals (NB100-690, 1: 1000).

### 2.6. Parental ssDNA assay

To visualize parental ssDNA in U2OS or FBH1 KO cells in response to Wee1i inhibition, 25,000 cells per sample were plated on circle coverslips in a 12-well plate and grown overnight. Cells were pretreated with 50 μM Idu for 48 hrs (Idu was refreshed after 24 hrs). Cells were then treated with 200 nM or 400 nM adovasterib or DMSO (vehicle control) for 8 or 24 hrs before cell fixation. Cells were stained using BrdU and PCNA antibodies following the immunofluorescence protocol.

### 2.7. Western Blotting

Cells were lysed (10^6^ cells/100 μL) in 1X SDS Buffer (062.5 mM Tris-HCL pH 6.8, 2% SDS, 10% glycerol, 0.1 M DTT, and 0.1 mg/mL bromophenol blue). Samples were run on a 10% SDS-PAGE gel and transferred to PVDF membrane. Membranes were blocked with 5% (wt/vol) powdered milk (Nestle Carnation) dissolved in TBST (10 mM Tris, 150 mM NaCl, 1% [vol/vol] Tween 20) for 1 hr at room temperature. Membranes were incubated overnight at 4°C in primary antibodies diluted in 5% milk in TBST. Membranes were washed in TBST for 30 min with 3 buffer changes. Blots were incubated with HRP-conjugated antibodies (Licor WesternSure, 1:2,000) for 1 hr at room temperature and washed as described above. Cells were incubated with WesternSure Chemiluminescent Substrate (Licor). Blots were imaged using the C-digit imaging system (Li-COR). Membranes were stripped for 30 min with 2 buffer changes with a mild stripping buffer (15 mg/mL glycine, 1 mg/mL SDS, 10% Tween 20, pH adjusted to 2.2 using concentrated HCL), washed with TBST before reprobing with antibodies.

### 2.8 TUNEL Assay

U2OS and FBH1 KO cells (50,000) were plated on coverslips in a 12-well plate. Cells were treated with 400 nM adovasterib or DMSO for 24 hrs. Cells were fixed as done for immunofluorescence. The TUNEL assay was performed using the Click-It Plus TUNEL Assay (Invitrogen) as per manufacturers’ instructions. Images were scored using Fiji software.

### 2.9. Annexin V staining

U2OS or FBH1 KO cells (4×10^5^) were seeded into 6 cm dishes and grown overnight. Cells were treated with either DMSO or 400nM Wee1i for 48 hrs. Culture media was collected and adherent cells were harvested by trypsinizing cell monolayer for 5 min at 37℃. Culture media and cells were collected by centrifugation at 1000 x g for 5 min. Cells were stained with Annexin V and propidium iodide using Annexin V apoptosis detection kit (Invitrogen Cat# 50-929-7) as per manufacturers’ instructions. Cells were analyzed by flow cytometry using a CytoFLEX (Beckman) and FCS express 7 software v7.16.0047 (De Novo Software).

### 2.10. Mitotic index

U2OS or FBH1 KO cells (4×10^5^) were seeded into 6 cm dishes and grown overnight. Cells were treated with either DMSO or 400 nM Wee1i for 8 hrs. Culture media was collected and adherent cells were harvested by trypsinizing cells for 5 min at 37℃. Cells were fixed by adding 5 mL of ice-cold 70% EtOH in 1X PBS (v/v) while vortexing. Samples were stored at - 20℃ until staining. Fixed cells were washed with 1X PBS and permeabilized with 0.25% Triton X-100 in 1X PBS (v/v) by incubating on ice for 15 min. Cells were washed with 1X PBS prior to incubation with anti-pH3 Ser10 antibody (Abcam, cat#ab5176, 1:1000) in 3% BSA in 1X PBS (w/v) overnight at 4℃. Cells were washed with 1X PBS and incubated with AlexaFluor 594 antibodies (1:250) for 30 min at room temperature. Cells were incubated in 3 µM DAPI (w/v in 1X PBS) for 15 min at room temperature. Cells were washed with 1X PBS and incubated with RNaseA (250µg/mL in 1X PBS (v/v)) for 30 min at 37℃. Samples were analyzed by flow cytometry using a CytoFlex (Beckman) and FCS express 7 software.

### 2.11. Statistical analysis

Values are presented as means ± standard deviation of at least three independent experiments unless otherwise indicated. Means of treated groups were compared against those of the appropriate control, and statistical analyses were performed using GraphPad Prism 9 (v9.0.0; Dotmatics). Statistical significance was determined using Analysis of Variance (ANOVA) followed by a Tukey HSD test. A *p*-value less than 0.05 was considered statistically significant.

## 3. RESULTS

### 3.1. Treatment of FBH1 KO cells with WEE1i increases genome instability

To determine how FBH1 impacts the response to WEE1 inhibition, we knocked out FBH1 in U2OS and confirmed loss of FBH1 protein by western blotting (Fig. 1A). We treated U2OS and FBH1 KO cells with the WEE1 inhibitor, adovasterib (hereafter WEE1i) [12]. Treatment of cells with WEE1i significantly reduced levels of CDK1 phosphorylation at Y15 indicating efficient inhibition of WEE1 (Fig 1B).

**Fig. 1.**
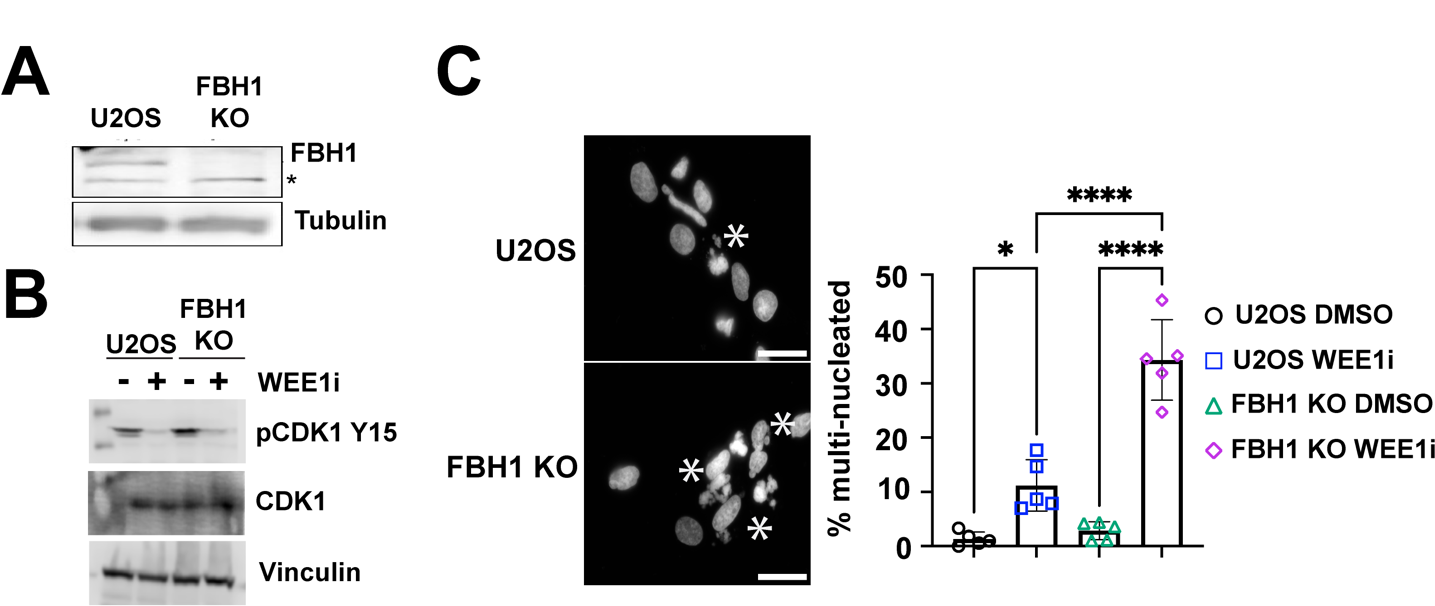
Loss of FBH1 in U2OS cells leads to increased multinucleation in response to Wee1 inhibition. (**A)** Representative Western blot images showing knockout of FBH1 in U2OS cells. Tubulin is used as a loading control. Asterisks indicate a non-specific band. (**B)** Representative western blot images showing levels of phospho-CDK at residue Y15 in U2OS or FBH1 KO cells treated with 400 nM WEE1i for 24 hrs. CDK1 and Vinculin are loading controls. (**C)** Representative images showing nuclear aberrations in U2OS or FBH1 KO cells treated 400nM WEE1i for 24 hrs. Scale bar= 10 μm. Asterisks indicate nuclei that were scored as multinucleated. Bar graphs represent the mean percentage of multi-nucleated cells (N=5). Error bars are standard deviation. * *p*<0.05, **** *p*<0.00005, ANOVA, Tukey HSD.

In cancer cells, forced mitotic entry in cells with DNA damage leads to mitotic catastrophe and cell death. We treated U2OS and FBH1 KO cells with WEE1i for 24 hrs and examined cells for nuclear abnormalities. Specifically, we determined if WEE1i treatment led to multi-nucleated cells indicative of mitotic catastrophe (Fig 1C). In U2OS cells, WEE1i significantly increased the percentage of multinucleated cells from 1.3% to 11.2%. Strikingly, FBH1 KO cells exhibited a 3-fold increase in multinucleated cells after WEE1i treatment (34.3% cells) compared to U2OS suggesting that FBH1 prevents mitotic catastrophe after WEE1 inhibition.

### 3.2. Reduced ssDNA accumulation in FBH1 KO cells after WEE1i treatment

In response to WEE1i, increased origin firing leads to exhaustion of nucleotide pools triggering replication fork stalling and accumulation of single-stranded DNA (ssDNA) [3]. Given previous reports showing FBH1 is required for the cellular response to hydroxyurea, we determined how FBH1 KO cells respond to WEE1i treatment by examining cells for markers of replication stress. First, we measured accumulation of ssDNA using the parental ssDNA assay [20]. U2OS and FBH1 KO cells were grown in the presence of IdU for 48 hrs before releasing into media with and without WEE1i. In U2OS cells, WEE1i treatment significantly increased the number of S phase (i.e., PCNA positive) cells containing IdU (24.5%) indicating WEE1i results in accumulation of ssDNA (Fig. 2A, B). Compared to U2OS cells, FBH1 KO cells had significantly fewer IdU positive cells (8.7%). We observed similar results when U2OS and FBH1 KO cells were treated with 200 nM (14.2% vs 4.1%, respectively; **Fig. S1A, B**) or when treated with 400 nM for 8 hrs (19.0% vs 8.1%, respectively; **Fig. S1C, D**). Together, these data indicate FBH1 promotes parental ssDNA accumulation in response to inhibition of WEE1.

**Fig. 2.**
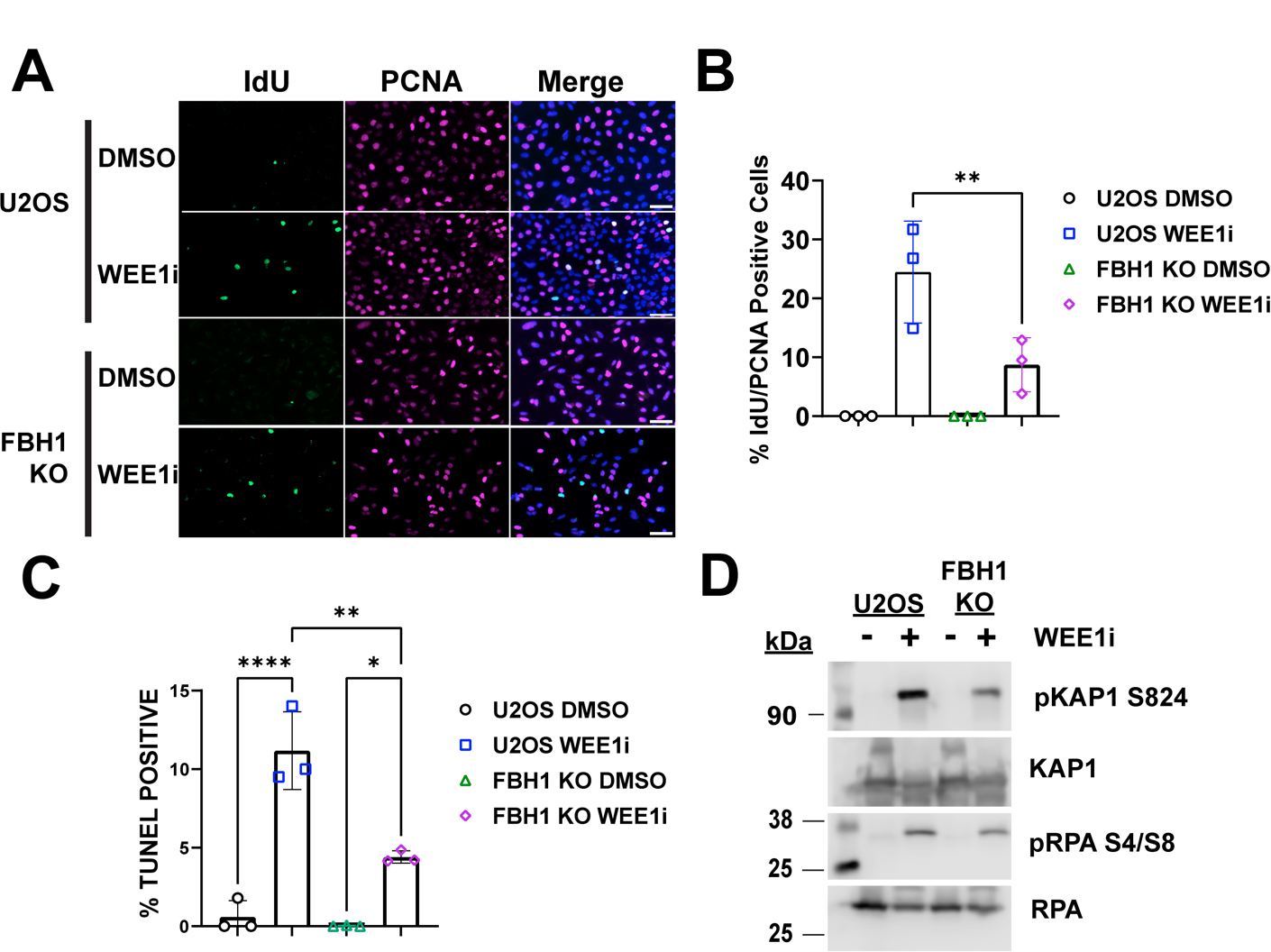
Impact of FBH1 KO on the replication stress response after treatment with WEE1i. (**A**) Representative images depicting ssDNA (IdU) in S phase cells (PCNA positive) in U2OS or FBH1 KO cells treated with 400 nM WEE1i for 24 hrs. Nuclei were counterstained with DAPI (blue). Scale bar= 60 μm. (**B**) Bar graph represents the mean percentage of PCNA positive cells containing ssDNA with individual data points (N=3). Error bars are standard deviation. (**C)** The mean percentage of TUNEL positive (**D)** Western blots depicting pKAP1 S824 and pRPA S4/8 levels after treatment with WEE1i for 24 hr. KAP1 and RPA were used as loading controls. * *p*<0.05, ** *p*<0.005, **** *p*<0.00005. ANOVA; Tukey HSD.

### 3.3. FBH1 is required for double strand break formation after WEE1i inhibition

WEE1 inhibition results in unrestrained cleavage by the endonuclease MUS81/SLX4 [21]. In response to hydroxyurea, FBH1 promotes MUS81-dependent cleavage of replication forks. To determine if FBH1 promotes double strand break formation after treatment with WEE1i, we performed the TUNEL assay that measures DNA breaks that occur due to DNA fragmentation during apoptosis or cleavage of replication forks due to replication catastrophe [6]. In U2OS cells, WEE1i treatment increased the percentage of TUNEL positive cells (18-fold) indicating formation of double strand breaks (Fig. 2C). In FBH1 KO cells treated with WEE1i, we observed a 2.5-fold reduction in TUNEL positive cells compared to U2OS indicating FBH1 promotes efficient DNA break formation.

After replication fork collapse, RPA undergoes phosphorylation at serine 4/8 (pRPA S4/8) and KAP1 undergoes phosphorylation at serine 824 (pKAP1 S824) by ATM and DNAPKcs [22,23]. We examined pKAP1 and pRPA induction by western blotting in U2OS and FBH1 KO cells after treatment with WEE1i (Fig. 2D). In U2OS cells, we observed a significant increase in pKAP1 and pRPA after WEE1i treatment consistent with double strand break formation due to replication fork collapse. Compared to U2OS cells, FBH1 KO had a significant reduction in pRPA S4/8 and pKAP1 S824 indicating reduced activation of double strand break signaling. Together, these data indicate FBH1 is required for the cellular response to replication stress induced by WEE1i treatment.

### 3.4. WEE1i treatment leads to two distinct pan-nuclear γH2AX populations in U2OS cells

The histone H2AX is phosphorylated at Ser139 in response to DNA damage to form immunostaining foci. In cells undergoing replication catastrophe or apoptosis, nuclei exhibit intense γH2AX staining known as pan-nuclear staining [6,24,25]. Previous studies have shown FBH1-deficient cells have a significant reduction in pan-nuclear γH2AX in response to replication stress caused by hydroxyurea treatment. Given the reduction in markers of replication catastrophe in FBH1 KO cells after treatment with WEE1i, we measured the induction of pan-nuclear γH2AX in U2OS and FBH1 KO cells after treatment with WEE1i (Fig. 3, **S2**). In U2OS cells, WEE1i treatment resulted in 30% of cells being positive for pan-nuclear γH2AX staining (**Fig. S2B**). We did not observe a significant difference in the percentage of cells with pan-nuclear γH2AX in FBH1 KO cells (33.7%). This result suggests that FBH1 is not required for γH2AX induction in response to WEE1i.

**Fig. 3.**
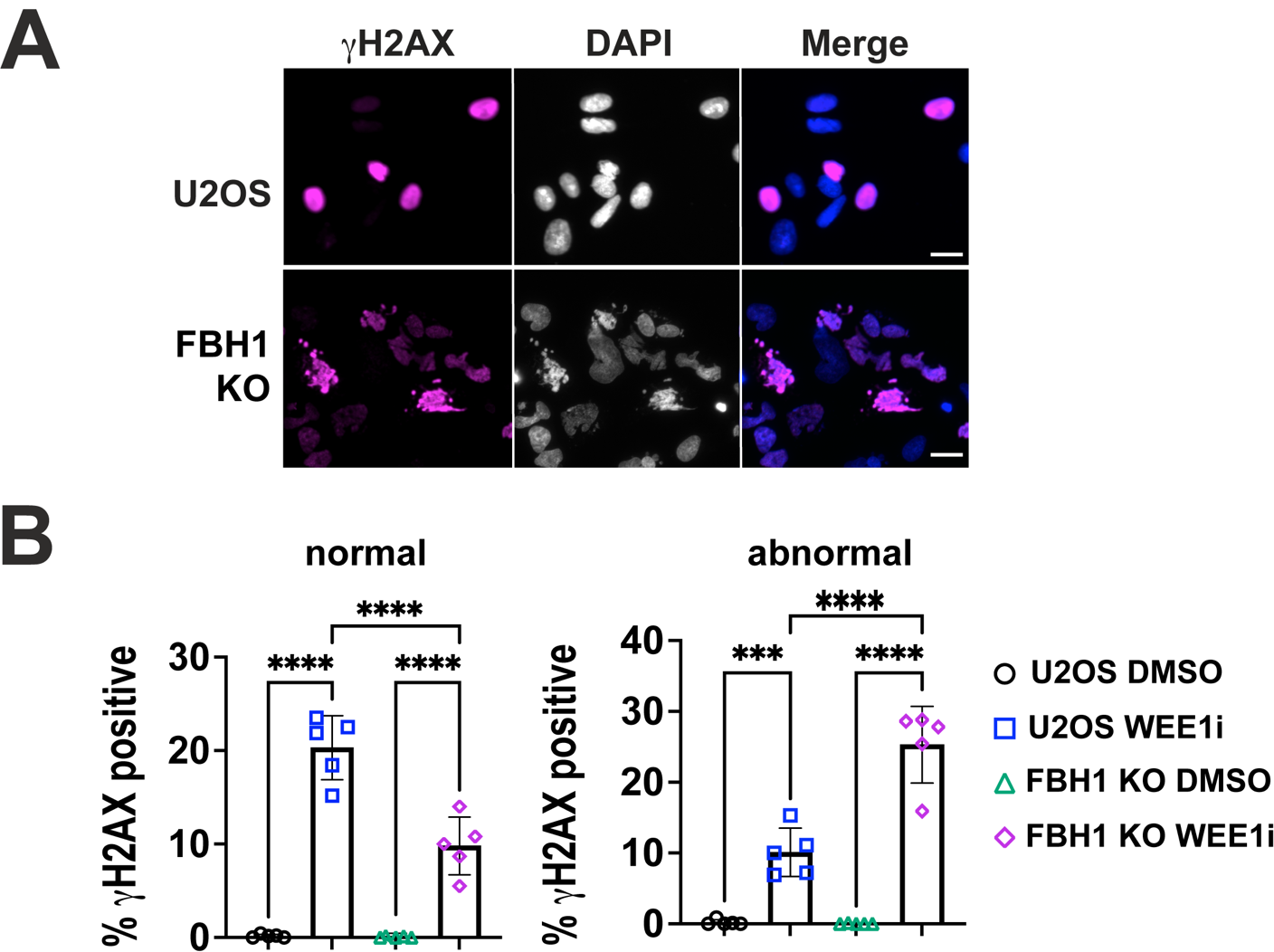
WEE1i treatment leads to two distinct pan-nuclear γH2AX populations in U2OS and FBH1 KO cells. **(A)** Representative images of U2OS or FBH1 KO cells treated with DMSO or 400nM WEE1i for 24 hrs. Cells stained with γH2AX (magenta). DNA was stained with DAPI. Scale bar=60 μm. (**B)** Bar graphs represent the mean percentage of pan-nuclear γH2AX positive cells in U2OS or FBH1 KO cells treated with DMSO or 400nM WEE1i for 24 hrs. Error bars are standard deviation. Left graph depicts γH2AX positive cells in cells without nuclear aberrations (normal). Right graph depicts γH2AX positive cells in abnormal nuclei. *** *p*<0.0005, **** *p*<0.00005. ANOVA, Tukey HSD.

Given the reduction in ssDNA accumulation and double strand break formation observed in FBH1 KO cells, we decided to investigate further why we did not observe a defect in pan-nuclear γH2AX formation in FBH1 KO after WEE1i treatment. We found the nuclei staining positive for pan-nuclear γH2AX differed between U2OS and FBH1 KO cells (Fig. 3A, B). Of the 30% γH2AX positive cells in U2OS cells, 20% (60% of γH2AX positive population) occurred in nuclei without any aberrations (i.e., normal). Of the 33.7% γH2AX positive nuclei in FBH1 KO cells, 25.3% (70.4% of γH2AX positive cells) occurred in multi-nucleated cells. This result suggests pan-nuclear γH2AX is being induced in distinct subpopulations of cells in U2OS and FBH1 KO cells.

### 3.5. Reduced mitotic entry in FBH1 KO cells treated with WEE1i

WEE1 inhibition results in premature mitotic entry resulting in mitotic catastrophe in cells with unrepaired DNA damage[9,13]. To determine if the increase in mitotic catastrophe observed in FBH1 KO cells was due to an increase in unscheduled mitotic entry, we treated cells with 400 nM WEE1i and examined phosphorylated histone H3 positive Ser 10 (pH3+) after 8 hrs by flow cytometry (Fig 4A**, S3**). In untreated controls, 2.15% of U2OS and 2.41% of FBH1 KO were pH3 positive. In response to WEE1i, we observe a 6.6-fold increase in pH3+ cells in U2OS and a 4-fold increase in FBH1-KO cells. Thus, FBH1-deficiency results in a signicantly lower percentage of pH3+ cells. Based on DNA content, it appears FBH1 KO cells may exhibit a higher percentage of cells undergoing premature mitotic entry (e.g., pH3+ cells in S phase, **Fig. S3**). However, the severe nuclear abnormalities observed in FBH1 KO cells precludes our ability to perform a cell cycle analysis after WEE1i treatment. Given the overall reduction in pH3+ cells observed in FBH1 KO cells after WEE1i treatment, the premature mitotic entry we observe in FBH1 KO cells cannot account for the 3-fold increase in miotic catastrophe we observe after WEE1i treatment.

**Fig. 4.**
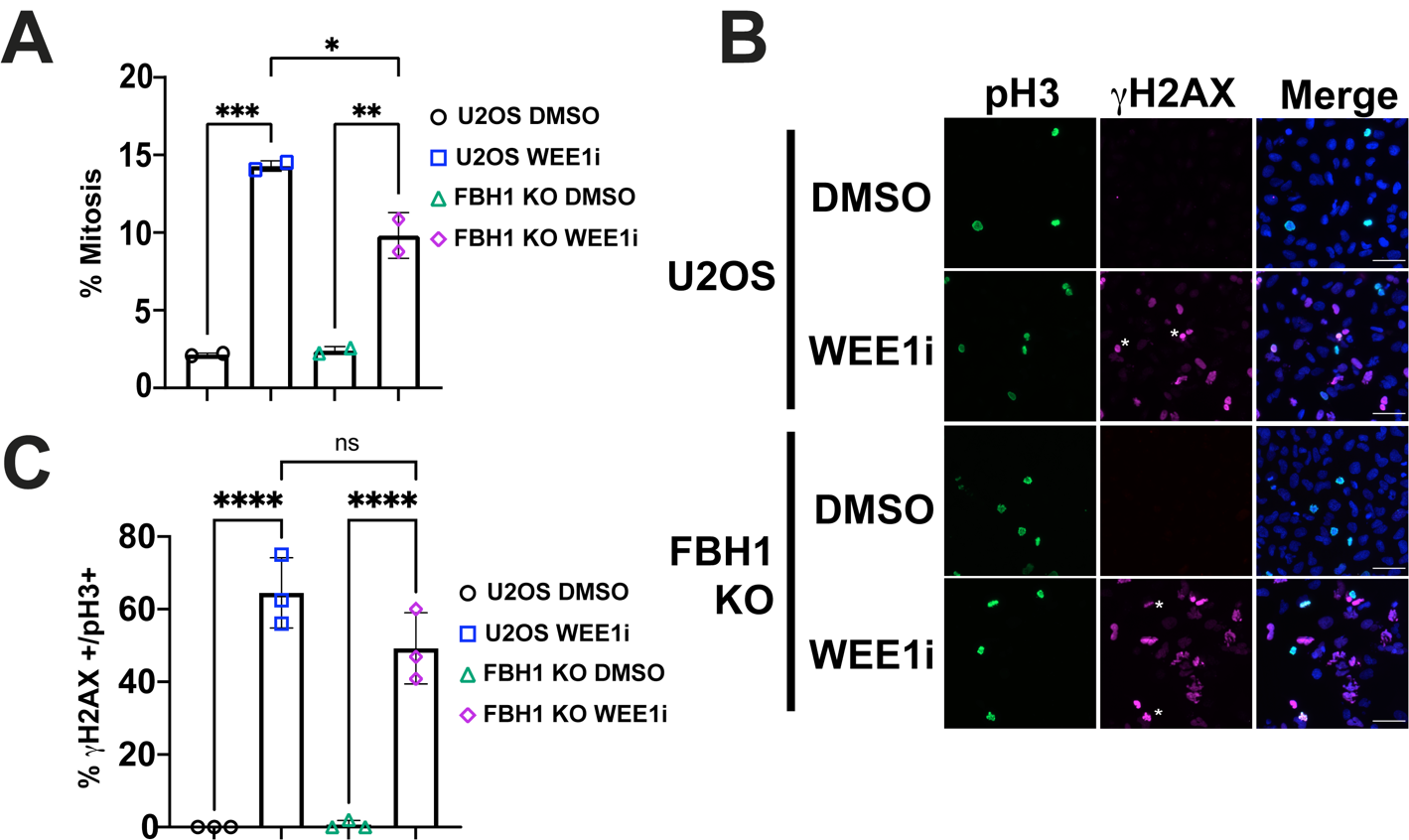
Mitotic DNA damage in FBH1 KO cells treated with WEE1i. **(A)** Bar graph depicts mean percentage of U2OS for FBH1 KO cells in mitosis (pH3 positive cells) after treatment with DMSO or 400 nM WEE1i for 8 hrs as determined by flow cytometry. Error bars are standard deviation. N=2. (**B**). Representative images of U2OS or FBH1 KO cells treated with 400 nM WEE1i for 24 hrs and stained for γH2AX (magenta), pH3 (green), and countered stained with DAPI (blue). Scale Bar= 60 μm. **(C)** Bar graph represents mean percentage of nuclei staining positive for γH2AX and pH3. Error bars are standard deviation. N=3. ns-not significant, * *p*<0.05, \*\**p*<0.005, *** *p*<0.0005, **** *p*<0.00005. ANOVA; Tukey HSD

To determine if FBH1 KO cells exhibited an increase in DNA damage in mitosis compared to U2OS, we examined pH3 positive cells for pan-nuclear γH2AX (Fig. 4B, C). In untreated U2OS or FBH1, we did not observe γH2AX staining in mitotic cells. After WEE1i treatment, 64.5% of pH3 positive cells stained positive for γH2AX in U2OS cells. In FBH1 KO cells, we observed a small, but insignificant decrease (49.5%) in pH3+ cells staining positive for γH2AX. This result indicated that loss of FBH1 does not increase DNA damage in mitotic cells. Together, these results indicate the increase in mitotic catastrophe in FBH1 KO cells is not due to an increase in mitotic entry or increase in mitotic DNA damage.

### 3.7. FBH1 KO cells exhibit reduced growth in response to WEE1i

Cell lethality caused by WEE1i treatment can occur through mitotic catastrophe or replication catastrophe[3,13,26]. In FBH1 KO cells, we observed an increase in mitotic catastrophe, but a decrease in replication catastrophe. To determine how FBH1 status impacted cell survival after WEE1i treatment, we measured apoptosis by staining cells with Annexin V after treatment with WEE1i for 24 hrs (Fig. 5A). We observed significant variation in Annexin V positive populations in response to WEE1i in both U2OS and FBH1 KO cells. However, on average we did not observe a significant difference in the percentage of Annexin V positive cells in U2OS (14.9%) and FBH1 KO (14%). Thus, FBH1 is not required to initiate apoptosis in response to WEE1 inhibition.

**Fig. 5.**
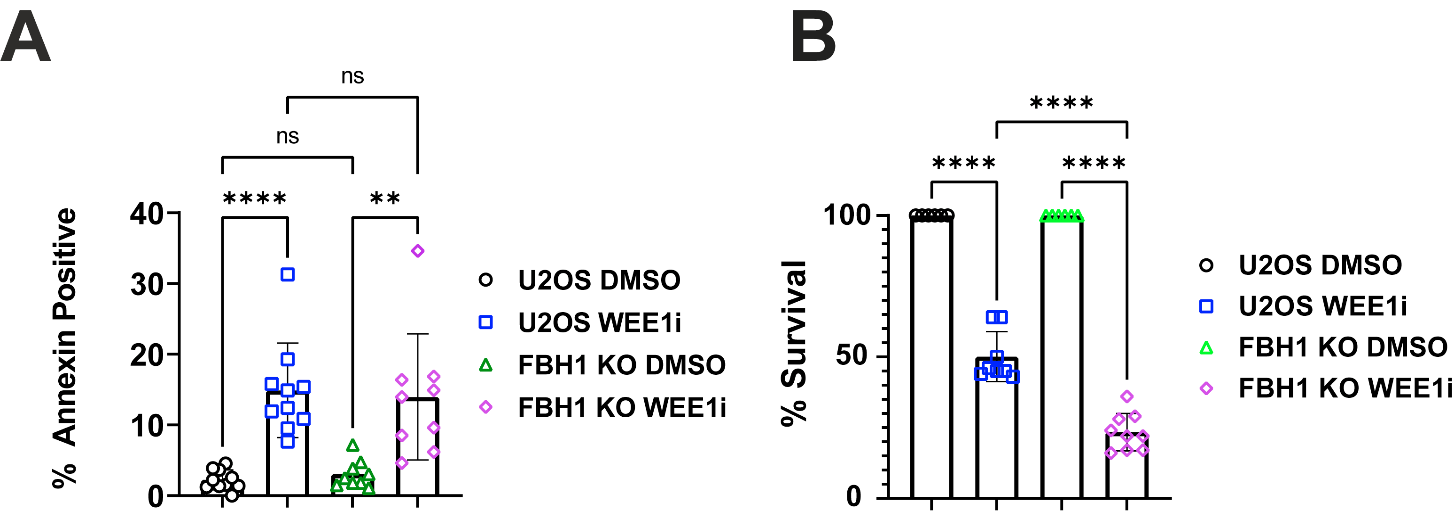
FBH1 KO cells are sensitive to WEE1 inhibitors. (**A**) Annexin V staining in U2OS and FBH1 KO cells were treated with DMSO or 400 nM WEE1i for 8 hrs. Bar graph represents the mean percentage of Annexin V positive cells of at least 3 separate trials. Error bars are standard deviation. (**B**) Cell survival of U2OS and FBH1 KO cells after treatment with DMSO or 400 nM WEE1i for 48 hrs. ns-not significant \*\**p*<0.005, *** *p*<0.0005, **** *p*<0.00005. ANOVA; Tukey HSD.

Next, we examined the ability of FBH1 KO cells to recover from WEE1i by measuring cell survival (Fig. 5B). In U2OS cells, WEE1i resulted in a significant reduction in cell growth (50% compared to untreated). Compared to U2OS, FBH1 KO cells had a 2-fold reduction in cell growth (23.4% compared to untreated). Together, these results indicated FBH1 KO cells are more sensitive to WEE1 inhibitors.

## 4. DISCUSSION

Our results demonstrate FBH1 is required to promote the cellular response to replication stress induced by WEE1 inhibition. Unscheduled origin firing in the absence of WEE1 results in nucleotide depletion and replication catastrophe, similar to the mechanism of action of the replication inhibiting drug, hydroxyurea[3]. FBH1 promotes MUS81-dependent fork breakage and double strand break signaling in response to replication stress induced by ultraviolet light and hydroxyurea[16–18]. Here, we observed a significant decrease in ssDNA accumulation, double strand break formation and double strand break signaling in FBH1-deficient cells treated with WEE1 inhibitors suggesting FBH1 plays a similar role in promoting breaks in response to replication stress induced by WEE1 inhibition.

Unlike hydroxyurea, the dampened replication stress response observed in FBH1-deficient cells is not resulting in increased resistance to WEE1 inhibitors. FBH1-deficient cells have reduced cellular survival compared to U2OS cells and induce apoptosis after WEE1i to similar levels as U2OS as determined by Annexin V staining. We performed the TUNEL assay and observed a decrease in TUNEL positive cells in response to WEE1i compared to U2OS cells. Although these results appear contradictory, TUNEL labeling detects DNA ends that arise during apoptosis and as a result of replication catastrophe whereas Annexin V staining is specific to apoptosis[6]. Thus, the reduction in TUNEL positive cells is likely due to a reduction in replication catastrophe in FBH1 KO cells after treatment with WEE1i. Given the increase in multinucleated cells that also stain positive for pan-nuclear γH2AX, we propose the reduction in cell survival observed in FBH1 KO cells is a result of mitotic catastrophe.

Why are FBH1-deficient cells more sensitive to WEE1i than U2OS cells despite a defect in the replication stress response? Cells with unrepaired DNA damage or under-replicated DNA must repair DNA damage before entering mitosis. WEE1 kinase plays a critical role in activation of the G2/M checkpoint in response to DNA damage[2]. We propose the sensitivity of FBH1 KO cells to WEE1 inhibition is due to the simultaneous increase in replication stress and loss of the G2/M checkpoint. In FBH1 KO cells, WEE1 inhibition could generate replication intermediates that sensitize cells to loss of the G2/M checkpoint. For instance, FBH1-mediated fork reversal could prevent PRIMPOL-mediated repriming at replication forks. In cells lacking fork reversal mediated by HTLF, SMARCAL1, and RAD51, up-regulation of PRIMPOL results in accumulation of ssDNA gaps[27,27]. Importantly, ssDNA gaps are filled during G2/M by post-replication pathways including translesion synthesis (TLS)[28]. Consistent with our proposed model, TLS-deficient cells are sensitized to WEE1 inhibitors[29]. Future work will determine how FBH1-mediated fork reversal inhibits pathways such as PRIMPOL-mediated repriming and TLS.

Our findings have important implications for the use of WEE1 inhibitors to target cancer cells. Hemizygous and homozygous loss of FBH1 was identified in 32% of cell lines derived from melanoma and this increased to 63% in cell lines derived from metastatic melanoma[18]. Our results suggest FBH1-deficient tumors may be sensitized to WEE1 inhibition. In addition, we found FBH1-deficient cells are sensitive to WEE1 inhibition despite having a significant reduction in replication catastrophe. The defect in the replication stress response in FBH1-deficient cells confers resistance to other replication stress inducing drugs including hydroxyurea. Thus, our results suggest that WEE1 inhibition may be a potential strategy to target tumors that are resistant to replication stress inducing drugs. We speculate the underlying replication defect in FBH1-deficient cells is ssDNA gap accumulation. Replication gap accumulation has been proposed to be a promising target for chemotherapy as defects in replication fork response have been shown to lead to gap accumulation in different genetic backgrounds including cells lacking BRCA1, BRCA2, and HLTF[27,29–31]. Further work will be needed to determine if gap accumulation is a potential biomarker to predict WEE1i sensitivity.

## Authorship contribution statement

J.M.M conceived the study. L.J, H.A.W, and J.M.M designed experiments, conducted experiments, analyzed the data, and interpreted the results. H.A.W. wrote the first draft of the manuscript. L.J., H.A.W, and J.M.M edited the manuscript

## Acknowledgements

This work was supported in part by funding from the American Cancer Society (RSG-21-175-01-DMC) and National Institutes of Health (R35 GM142512-01) to JMM. Equipment used in this study was purchased in part using funding from COBRE P20GM109094.

## FOOTNOTES

1 Abbreviations used are: KO, knockout; UV, ultraviolet; WEE1i, WEE1 inhibitors (adovasterib); ssDNA, single-stranded DNA; TLS, translesion synthesis

