## Supplemental Material for "FBH1 deficiency sensitizes cells to WEE1 inhibition by promoting mitotic catastrophe"

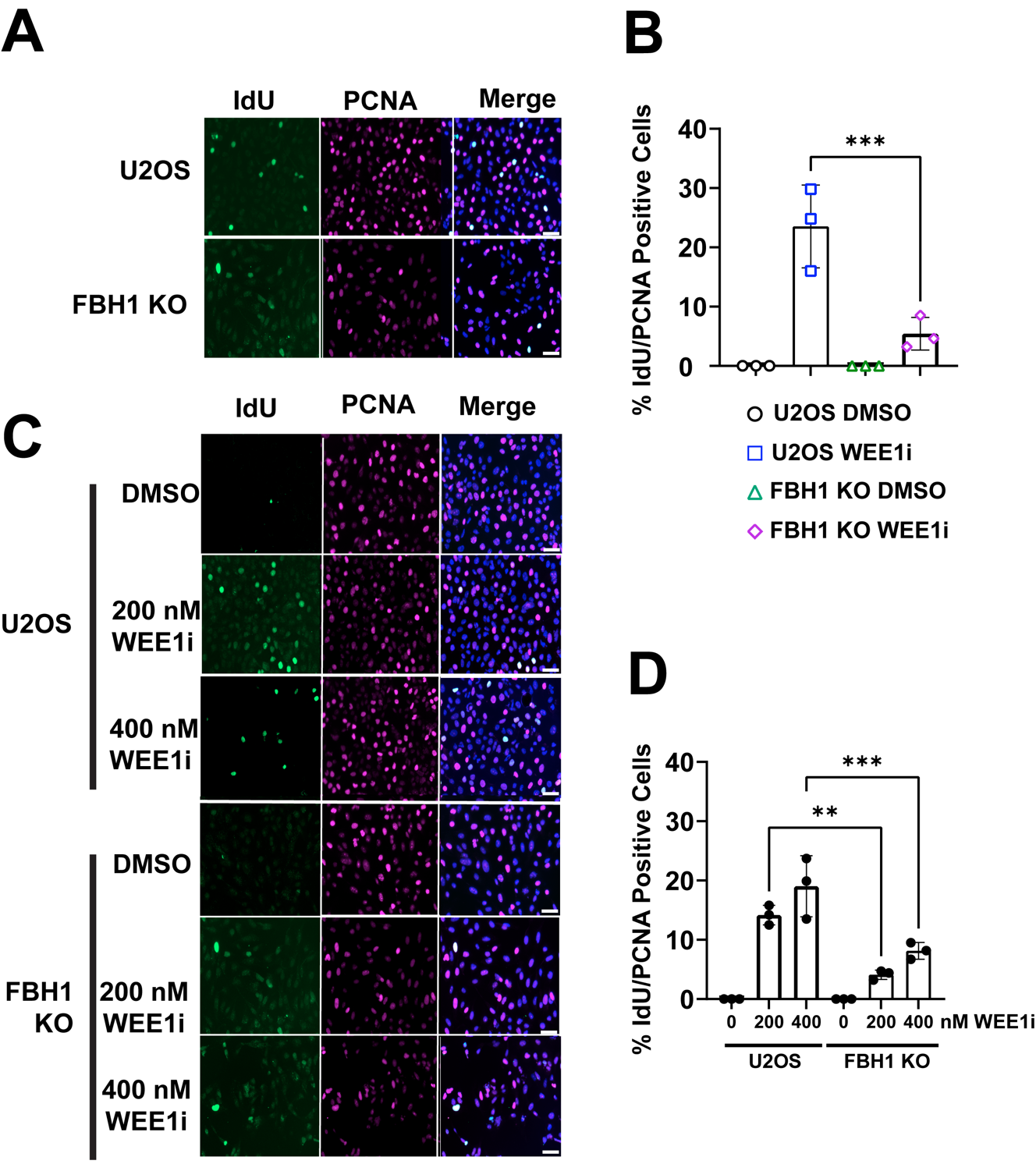

**Fig. S1. Reduced ssDNA accumulation in FBH1 KO cells treated with WEE1i.** (**A**) Representative images of U2OS or FBH1 KO depicting ssDNA in U2OS and FBH1 KO cells after the treatment with 200 nM WEE1i for 24 hrs. S phase cells were identified by staining with PCNA (red). Cells were stained with BrdU (IdU) to label exposed ssDNA (green). Nuclei were counterstained with DAPI (blue). Scale bar= 60 µm. (**B**) Bar graph represents mean percentage of PCNA positive cells containing ssDNA (IdU). Error bars are standard deviation. N=3. (**C)** Representative images of U2OS or FBH1 KO treated with DMSO, 200 nM or 400 nM WEE1i for 8 hrs. Cells were stained as in (A). Scale bar= 60 µm. (**D**) Bar graph represents mean percentage of PCNA positive cells containing ssDNA (IdU). Error bars are standard deviation. N=3. ** *p*<0.005, *** *p*<0.0005, ANOVA, Tukey HSD.

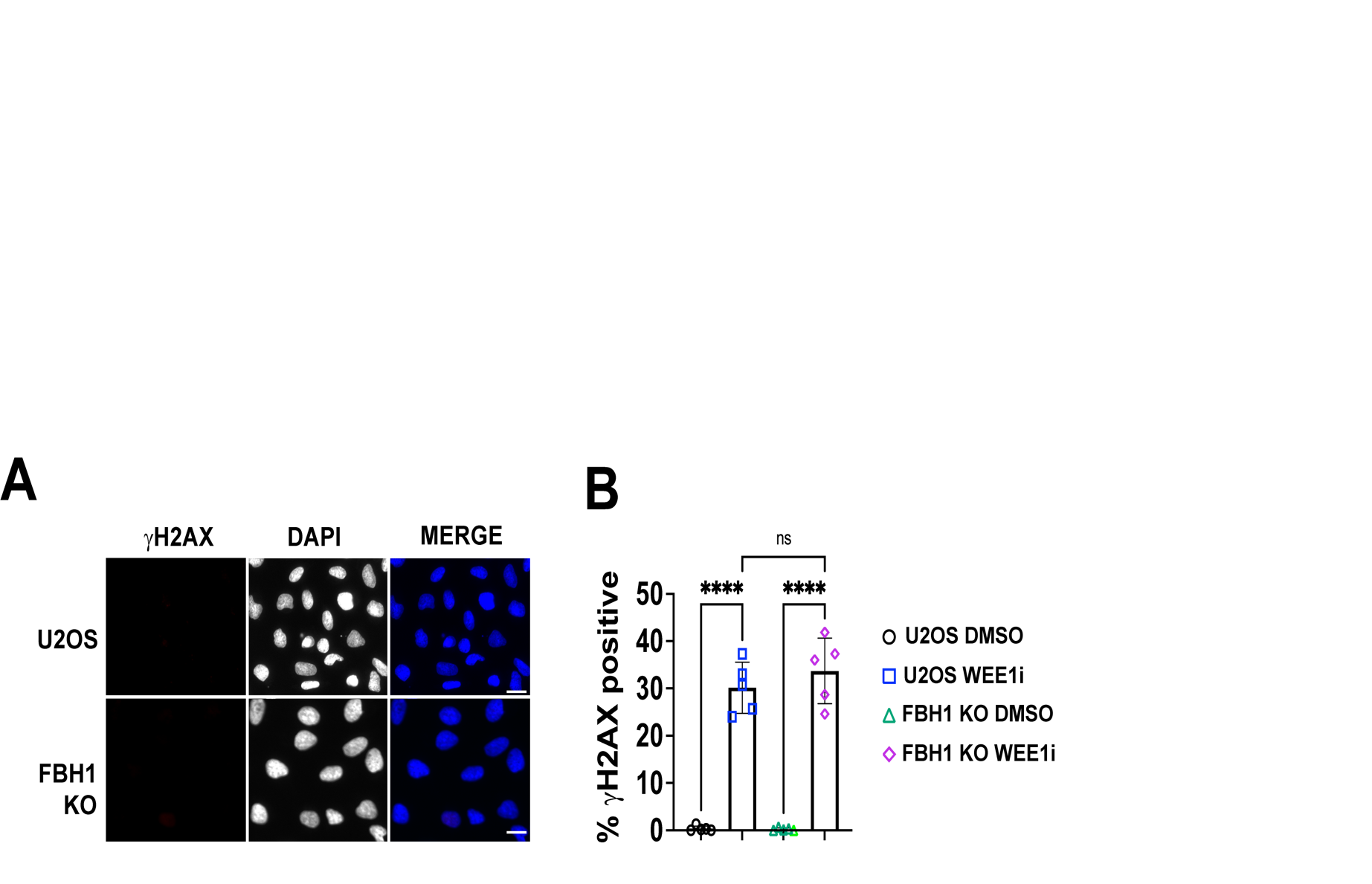

**Fig. S2. γH2AX induction in FBH1 KO cells after WEE1i treatment.** (**A**) Representative images of U2OS or FBH1 KO cells treated with DMSO with γH2AX (magenta) or DAPI (grayscale or blue). Control images for experiment in Fig. 3. Scale bar= 60 µm (**B)** Bar graph represents percentage of pan-nuclear γH2AX positive cells in U2OS or FBH1 KO cells, treated with DMSO or 400 nM WEE1i for 24 hours. Error bars are standard deviation. N=5. ns- not significant, **** p<0.00005, ANOVA, Tukey HSD.

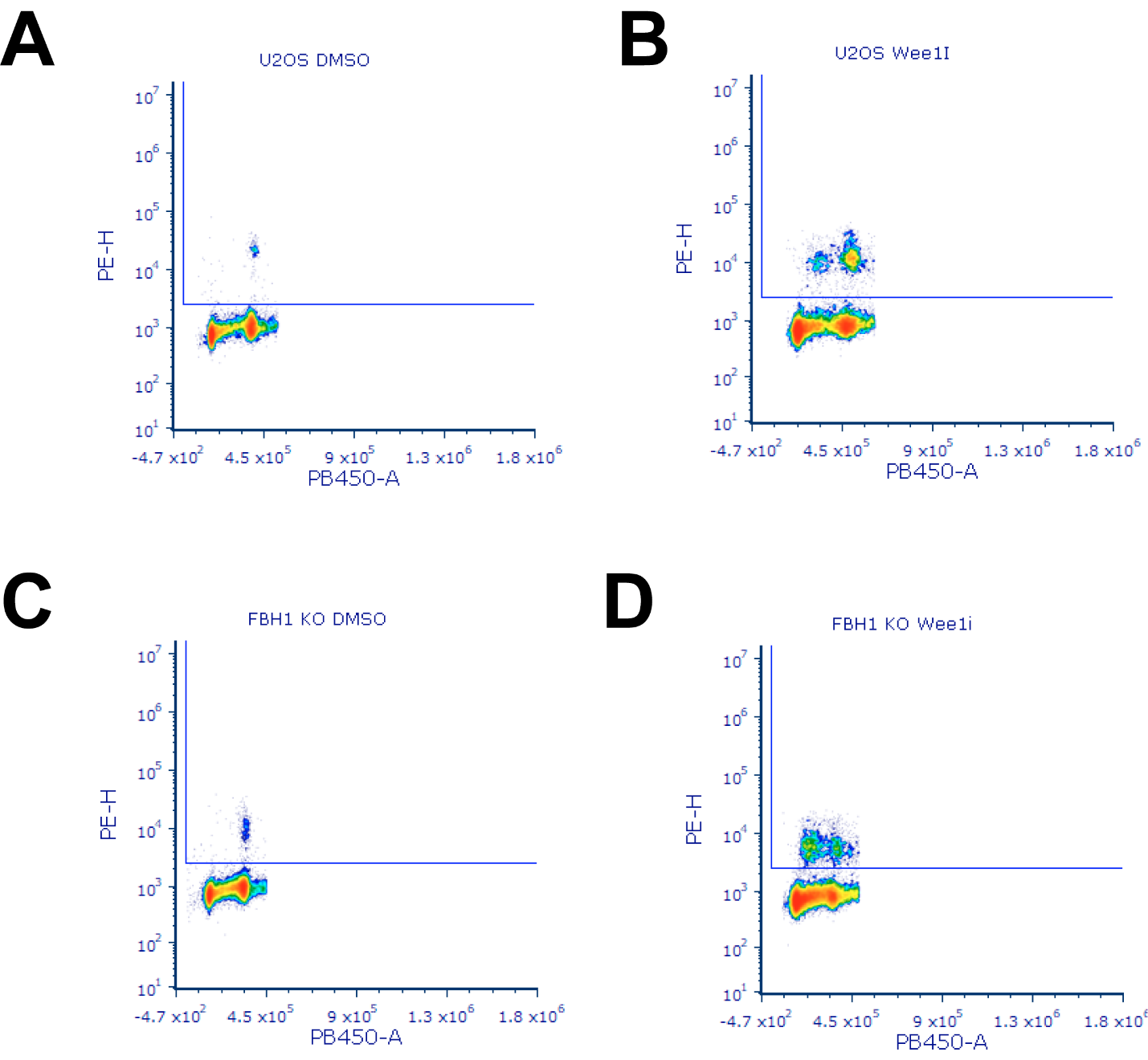

**Fig. S3. Mitotic cells in U2OS and FBH1 KO cells after treatment with WEE1i.** Representative dot plots of U2OS (**A&B**) or FBH1 KO cells (**C&D**) that were treated with DMSO or 400 nM WEE1i for 8 hours. Cells were stained with pH3 (PE-H) to label cells in mitosis and DAPI (PB450-A) to label DNA.
